## Supplementary figures and images for "Sex Differences in Discrimination Behavior and Orbitofrontal Engagement During Context-Gated Reward Prediction"

### Fig. S1

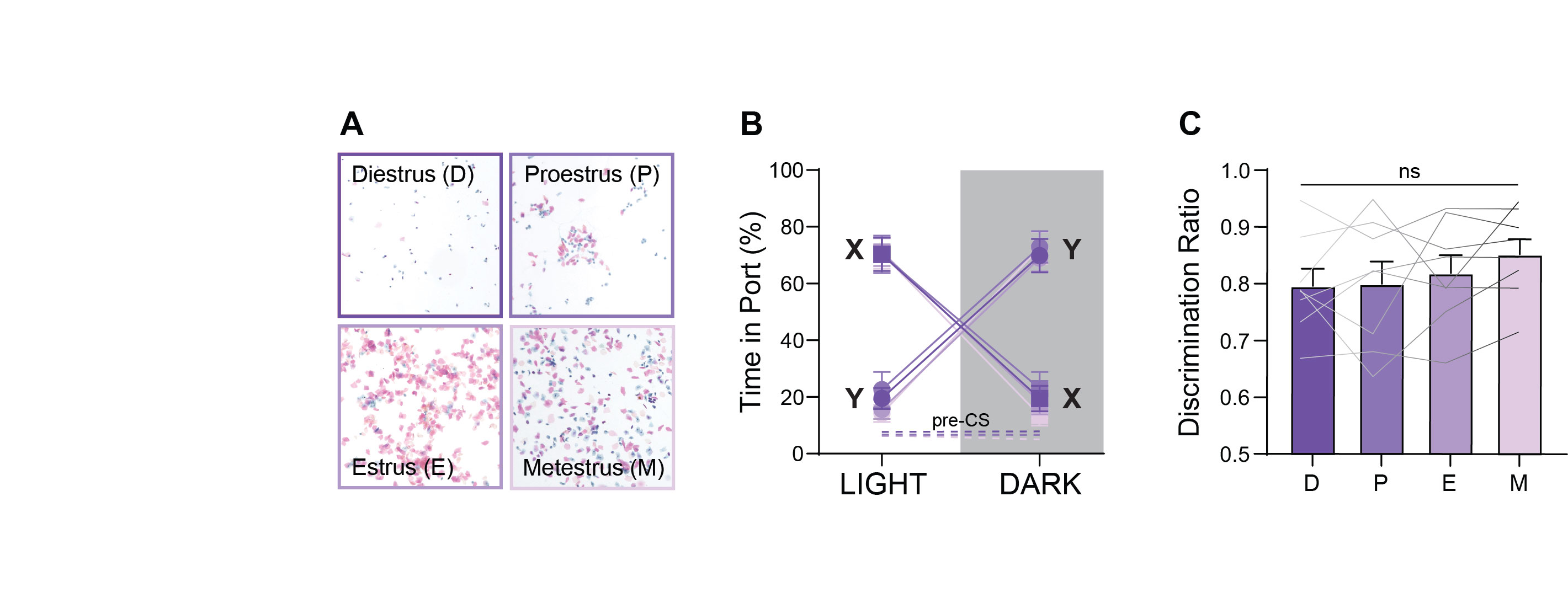
